# An adipocyte-specific lncRAP2 – Igf2bp2 complex enhances adipogenesis and energy expenditure by stabilizing target mRNAs

**DOI:** 10.1101/2020.09.29.318980

**Authors:** Juan R. Alvarez-Dominguez, Sally Winther, Jacob B. Hansen, Harvey F. Lodish, Marko Knoll

## Abstract

lncRAP2 is a conserved cytoplasmic adipocyte-specific lncRNA required for adipogenesis. Using hybridization-based purification combined with *in vivo* interactome analyses, we show that lncRAP2 forms ribonucleoprotein complexes with several mRNA stability and translation modulators, among them Igf2bp2. Transcriptome-wide identification of Igf2bp2 client mRNAs in white adipocytes reveals selective binding to mRNAs encoding adipogenic effectors and regulators. Depleting either lncRAP2 or Igf2bp coordinately downregulates these same target proteins. Ribosome profiling and quantitative proteomics show that this occurs predominantly at the level of mRNA, as binding of the lncRAP2-Igf2bp complex does not affect mRNA translation. Suppressing lncRAP2 or Igf2bp2 selectively destabilizes many mRNAs encoding proteins essential for energy expenditure, including Adiponectin, reducing adipocyte lipolytic capacity. Genome-wide association studies reveal specific association of genetic variants within both lncRAP2 and Igf2bp2 with body mass and type 2 diabetes, and we find that adipose lncRAP2 and Igf2bp2 are suppressed during obesity and diabetes progression. Thus, the lncRAP2-Igf2bp complex potentiates adipose development and energy expenditure and is associated with susceptibility to obesity-linked diabetes.

## Introduction

Pervasive transcription of the human and mouse genomes generates thousands of long noncoding RNAs (lncRNAs), yet only a minority have been linked to specific biochemical functions. Previous studies revealed that many lncRNAs are expressed only in white or/and brown adipocytes and play vital roles in adipocyte biology^1,2^, including adipogenesis, thermogenesis, and insulin sensitivity. We and others showed that the brown fat-specific lncRNAs Blnc1^3^ and lncBATE1^4^ interact with the nuclear matrix factor hnRNPU to mediate *trans*-activation of genes mediating the brown/beige thermogenic programs.

While lncRNAs can bind potentially DNA, RNA, or protein targets, much work suggests that lncRNA mechanisms predominantly involve binding to proteins, either as scaffolds of ribonucleoprotein complexes or as decoys to prevent their assembly^5,6^. To better understand how lncRNAs function, several technologies have been recently developed for unbiased mapping of lncRNA localization, protein targets, and functional domains^7,8^. Single molecule fluorescence in situ hybridization (smFISH) visually determines RNA abundance and location, showing whether a lncRNA diffuses to *trans* sites beyond its chromosomal locus or remains tethered *in cis* in the nucleus^9,10^. Cross-linking intact cells followed by hybridization-based RNA purification captures the specific DNA, RNA, or protein targets to which a lncRNA binds *in vivo* (rather than in solution after cell lysis)^11–14^. And footprint profiling can map the sites where protein complexes bind lncRNAs, revealing their functional domains^15–17^.

Here, we use these and other tools to interrogate the mode of action of lncRAP2, a white adipocyte-selective RNA essential for adipogenesis^18^. We show that lncRAP2 predominantly resides in the cytoplasm, yet it does not associate with ribosomes or other RNAs. Instead, lncRAP2 forms a complex with several RNA-binding proteins that affect mRNA stability and translation. Among these is Igf2bp2, which has been implicated in posttranscriptional control of metabolically-important proteins^19–21^, and we identify key adipogenic effectors and mediators of energy metabolism among its mRNA clients in white adipocytes. Indeed, we show that depleting either lncRAP2 or Igf2bp coordinately downregulates the proteins encoded by these targets, including Adiponectin, and that this occurs primarily through mRNA destabilization. Consistent with these findings, adipocytes in which either lncRAP2 or Igf2bp are depleted show compromised energy expenditure. Similarly, both in mice and humans the levels of lncRAP2 and Igf2bp2 in adipocytes are reduced during the development of obesity and diabetes. Analysis of genome-wide association studies further reveal a specific association of genetic variants within lncRAP2 and Igf2bp2 with increased body fat and greater risk of type 2 diabetes. Thus, a previously uncharacterized lncRAP2-Igf2bp2 complex regulates adipose energy expenditure, with implications for the susceptibility to and pathogenesis of obesity-linked diabetes in humans.

## Results

### lncRAP2 is a conserved adipocyte-specific cytoplasmic lncRNA required for adipogenesis

We previously identified several lncRNAs common to white and brown fat that, based on effects of depleting them in mouse preadipocytes, are essential for adipocyte development and function^18^. One, lncRAP2 (Genbank: NR_040299.1) is transcribed from an intergenic promoter bound by PPARγ, C/EBPα, and C/EBPβ (Fig. 1a and Supplementary Fig. 1a). 5’/3’ RACE verifies a capped and polyadenylated, 7.2kb spliced RNA (Supplementary Fig. 1b). lncRAP2 is highly adipose tissue-specific and strongly induced during early white and brown adipogenesis (Fig. 1c). Notably, lncRAP2’s structure, regulation, and expression traits are conserved in humans (Fig. 1b, d). Depleting lncRAP2 by ~70-75% by transducing Dicer-substrate siRNAs (DsiRNAs) into cultured primary white pre-adipocytes^22^ dramatically blocks adipogenesis, as evidenced by impaired lipid accumulation and reduced induction of key adipocyte genes (PPARγ, C/EBPα, Adiponectin, Fabp4, and Glut4) (Fig. 1e, f). Single molecule fluorescence in situ hybridization (smFISH) shows that lncRAP2 diffuses from the nucleus and spreads throughout the cytoplasm at 14 ±3 transcripts per white and 9 ±3 transcripts per brown adipocyte (Fig. 1h). Cell fractionation verifies its cytoplasmic localization, unlike lncRAP1 (also called FIRRE) (Fig. 1g), which mediates nuclear *trans*-chromosomal interactions among adipogenic genes^23^. Thus, lncRAP2 is a conserved cytoplasmic RNA essential for adipogenesis.

**Fig. 1.**
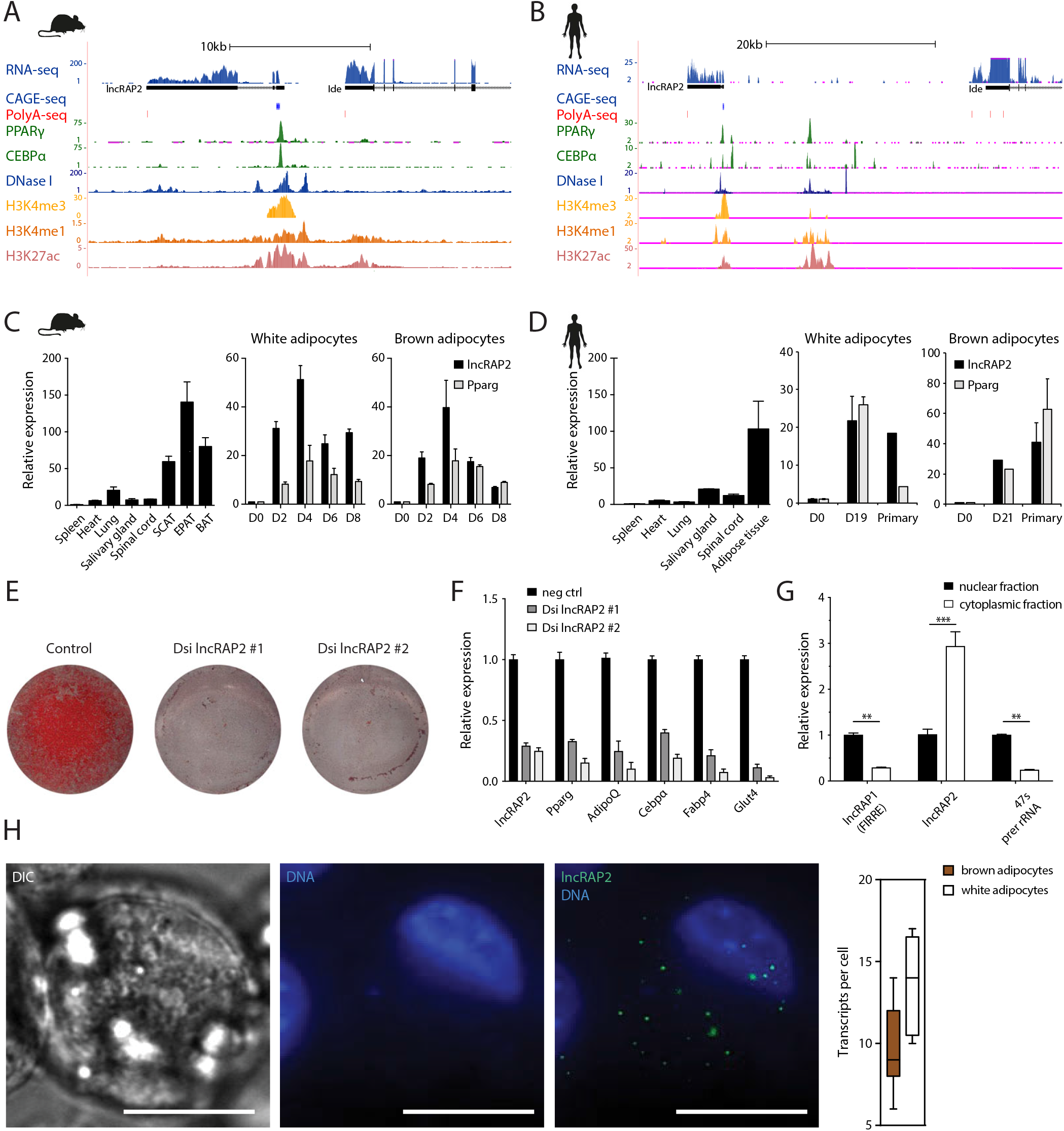
lncRAP2 is a conserved cytoplasmic RNA required for adipogenesis. (**a, b**) lncRAP2 is a capped, polyadenylated, and spliced RNA transcribed from a promoter bound by PPARγ, and C/EBPα. Tracks show signal from sequencing studies of white adipocytes from mouse (a) and human (b) (Supplementary Data 4) (**c, d**) lncRAP2 is adipocyte-specific and strongly induced during adipogenesis. Shown are relative tissue expression (normalized to spleen, left) and relative induction during *in vitro* differentiation of white (center) or brown (right) adipocytes from mouse (c) and human^71^ (d). SCAT, subcutaneous adipose tissue; EPAT, epididymal adipose tissue; BAT, brown adipose tissue. (**e, f**) lncRAP2 depletion blocks adipogenesis. Shown are lipid accumulation by Oil-red O staining (e) and relative expression of key adipocyte genes (f) in day 6 differentiated white adipocytes pre-treated with control or two different lncRAP2-targeting DsiRNAs. (**g, h**) lncRAP2 localizes to the cytoplasm. Relative expression in fractionated nuclear and cytoplasmic compartments (g), and single molecule FISH detection (h), in day 6 differentiated white adipocytes. lncRAP2 molecules above nucleus overlay DAPI staining in maximum z-stack projections of FISH images, quantified to the right (n=9 cells with ≥1 transcripts). **p <0.01, ***p <0.001 (t-test).

### lncRAP2 forms complexes with several proteins that regulate mRNA stability and translation, including Igfbp2

To investigate how lncRAP2 functions, we sought to identify its binding partners and targets. lncRAP2’s mRNA-like features suggest it could bind ribosomes, and to test this, we examined parallel RNA and ribosome footprint profiling during mouse adipogenesis^24^. Despite strong induction during adipogenesis, lncRAP2 is largely devoid of ribosome footprints and shows poor translatability (Fig. 2a, b). Supporting this, no peptides could be found in proteome surveys of brown or white adipocytes from either mouse or human^4,25^. Thus, lncRAP2 is unlikely to engage translating ribosomes.

**Fig. 2.**
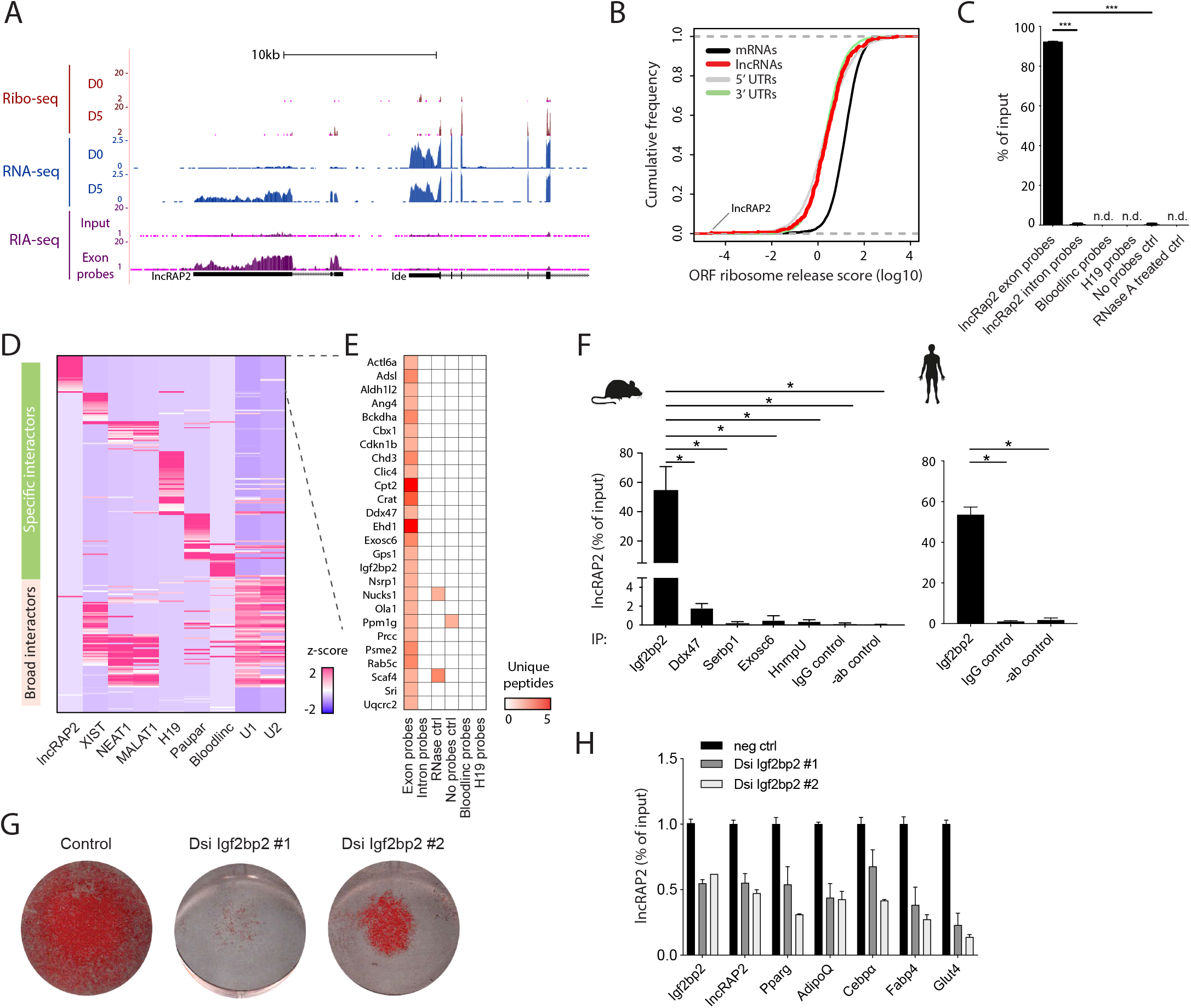
lncRAP2 forms a complex with mRNA stability and translation regulators. (**a, b**) lncRAP2 does not engage translating ribosomes. Tracks show signal from ribosome profiling, RNA, and RNA interactome sequencing studies of differentiated white adipocytes (a). Data are pooled from n=2-3 replicates. Translatability, measured by ribosome release from open reading frames after encountering a stop codon, is quantified in (b). (**c**) Efficient and specific enrichment of mature lncRAP2 by hybridization-based purification. Relative recovery of input lncRAP2 from differentiated 3T3-L1 adipocytes. ***p <0.001 (t-test). (**d, e**) ChIRP-MS identifies specific lncRAP2-binding proteins. The relative enrichment of high-confidence interactors captured by antisense purification of lncRAP2 in white adipocytes or other lncRNAs in other formaldehyde cross-linked cells is shown in (d). Unique peptide counts for specific lncRAP2 interactors are shown in (e). (**f**) Validation of lncRAP2 and Igf2bp2 direct interaction. Relative lncRAP2 recovery by native immunoprecipitates in white adipocytes from mouse (left) and human (right). *p <0.05 (t-test). (**g, h**) Igf2bp2 depletion blocks adipogenesis. Shown are lipid accumulation (g) and relative expression of key adipocyte genes (h) in day 6 differentiated white adipocytes pre-treated with control or Igf2bp2-targeting DsiRNAs.

To identify lncRAP2’s binding targets, we cross-linked intact mouse white adipocytes and used biotin-labeled smFISH probes to purify about 90% of cellular lncRAP2 (Fig. 2c). Less than 1% of cellular lncRAP2 was retrieved following RNase A treatment, without targeting probes, or with probes targeting introns or unrelated lncRNAs such as Bloodlinc (erythrocyte-specific)^26^ or H19 (a broadly expressed lncRNA)^27^, controls attesting to the efficiency and specificity of our RNA antisense purification protocol. We used RNA interactome analysis by sequencing (RIA-seq)^28^ to probe interactions of lncRAP2 with other RNAs *via* glutaraldehyde cross-linking. lncRAP2 was robustly enriched by distinct pools of exon-targeting probes (Supplementary Fig. 2a, b), but no other RNAs were cross-linked with significant enrichment or concordance between probe pools, indicating that lncRAP2 does not directly bind other RNAs.

We next conducted a comprehensive identification of RNA-binding proteins by mass spectrometry (ChIRP-MS)^14^ in 3T3-L1 adipocytes using formaldehyde cross-linking, which preserves both direct and indirect RNA-protein interactions. As controls, we compared crosslinked proteins to those captured using RNase A treatment, intron-targeting probes, non-targeting probes, or no probes. lncRAP2 purifications enriched protein analytes relative to these controls (Supplementary Fig. 2c), capturing 621 proteins with at least 2 unique peptides. We found 29 of these to be high-confidence (>2-fold enriched) lncRAP2-binding proteins (Supplementary Data 1). To identify specific lncRAP2 interactors, we compared these proteins to those captured by the hybridization-based purification of other lncRNAs: Xist, Neat1, Malat1, H19, Paupar, and Bloodlinc^14,29–32^ in formaldehyde cross-linked cells. Notably, 26 (90%) of the 29 interactors are lncRAP2-specific (Fig. 2d, e), and include regulators of fatty acid and keto acid metabolism (Crat, Cpt2, Bckdha) as well as mRNA translation and decay (e.g. Igf2bp2, Exosc6, Ddx47).

Among lncRAP2 interactors, Igf2bp2 regulates adipocyte function *via* posttranscriptional control of metabolically-important proteins, impacting sensitivity to diet-induced obesity and type 2 diabetes risk^19–21^. Native immunoprecipitation of endogenous Igf2bp2 in mouse and human white adipocytes specifically captures over 50% of lncRAP2 (Fig. 2f and Supplementary Fig. 2d), verifying direct lncRAP2-Igf2bp2 interactions. Igf2bp2 forms ribonucleoprotein complexes with client mRNAs to modulate their stability and translation^33–35^. Our data indicate that such complexes include Ddx47, with putative RNA unwinding activity in mRNA translation and decay^36^, and Exosc6/Mtr3, which binds to and presents mRNAs for degradation by the exosome^37^. Native immunoprecipitation of Ddx47 or Exosc6, however, captures less than 2% of cellular lncRAP2 (Fig. 2f), suggesting an indirect association to lncRAP2 *via* protein intermediates. We thus focused on the lncRAP2-Igfb2p connection.

### lncRAP2 and Igf2bp2 stabilize target mRNAs encoding metabolic effectors to potentiate adipocyte energy expenditure

Igf2bp2 is adipose tissue-enriched and like lncRAP2 is most highly expressed in white adipocytes (Supplementary Fig. 2e). We find little Igf2bp2 induction with adipogenesis, however, as it is already present in pre-adipocytes (Supplementary Fig. 2f, g). Depleting *Igf2bp2* in white pre-adipocytes by ~50% with DsiRNAs blocks their subsequent differentiation, analogous to lncRAP2 inhibition, reducing induction of adipogenic markers and lipogenesis (Fig. 2g, h). Induction of lncRAP2 is also disrupted, consistent with a differentiation block.

To investigate how Igf2bp2 regulates adipogenesis, we sought to identify its client RNAs. Unbiased sequencing of RNAs captured by native Igf2bp2-specific immunoprecipitates reveals 1,657 substrates enriched over input RNA or control purifications with IgG or no antibody (Supplementary Fig. 3a, Data 2). These comprise ncRNAs and mRNAs linked to various functions, with selective enrichment for adipogenic effectors (e.g. Nfat and Elovl proteins) and regulators (e.g. Cbp, Cited2, Ebf and Klf factors) (Fig. 3a). In line with known Igf2bp2 binding bias^33,38,39^, >60% of targets in adipocytes harbor CA-rich binding motifs, mainly enriched in 3’UTRs, and ~75% also copurify with Igf2bp2 in human cell types cross-linked with UV^40^ or 4-thiorudine^33^ (Supplementary Fig. 3b, c). Notably, substrates include mRNAs encoding key controllers of energy metabolism, such as the namesake Igf1/2 targets and PPARγ coactivator 1α (PGC1α) (Fig. 3b).

**Fig. 3.**
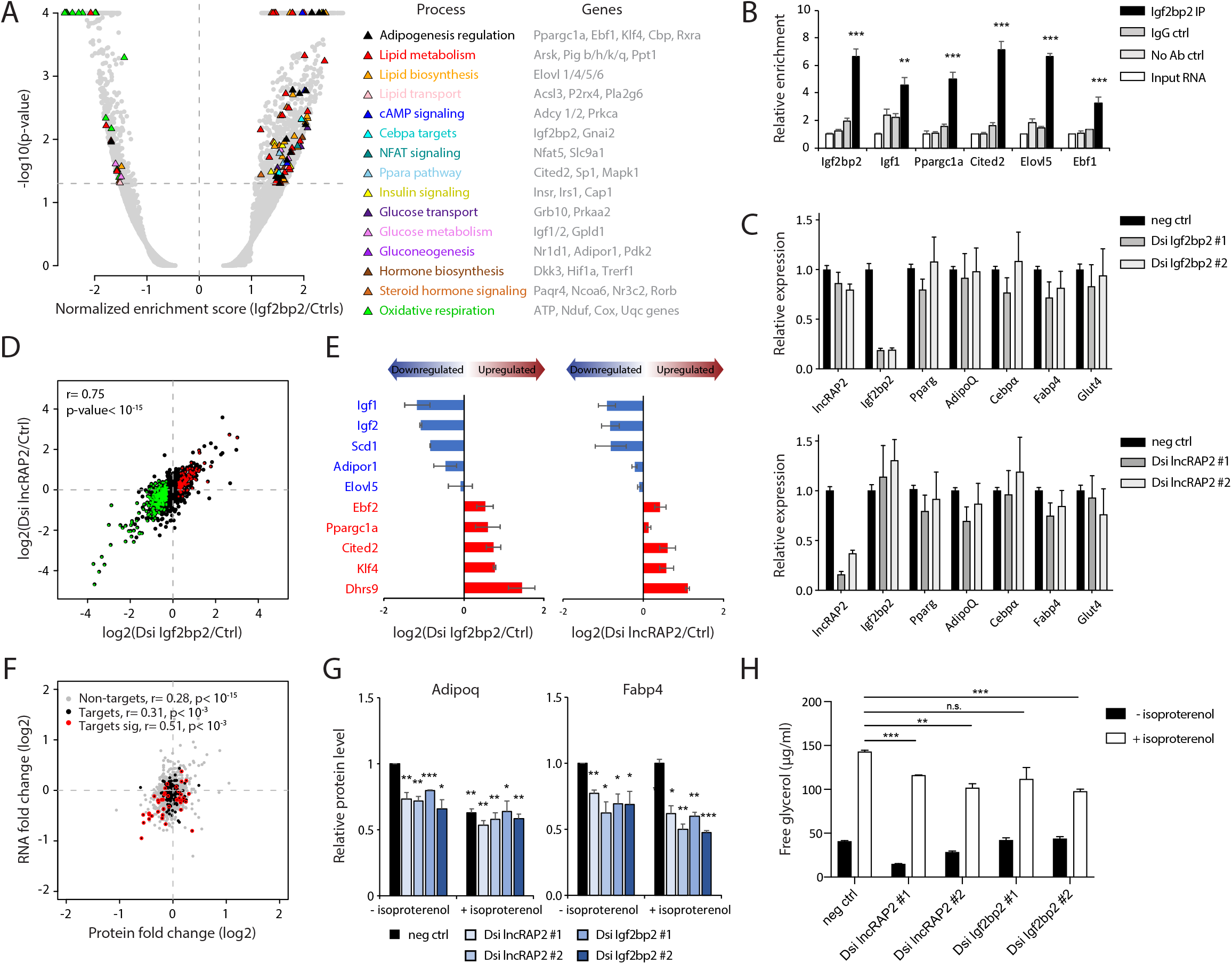
lncRAP2-Igf2bp2 complex regulates expression of target mRNAs encoding metabolically important proteins to potentiate adipocyte energy expenditure. (**a, b**) Igf2bp2 selectively binds mRNAs encoding adipogenic regulator and effector proteins in mouse white adipocytes. Shown is a gene set enrichment analysis highlighting significantly enriched (p <0.05) gene sets and their associated biological processes (a), with member genes shown to the right, and specific enrichment of mRNAs encoding key energy metabolism controllers (b), from native Igf2bp2 immunoprecipitations. **p <0.01, ***p <0.001 (t-test). (**c**) Depleting lncRAP2 or *Igf2bp2* in mature adipocytes does not alter expression of adipose marker genes or each other’s RNA levels. Relative expression of key adipocyte genes in day 6 differentiated white adipocytes pre-treated at differentiation day 4 with control, Igf2bp2-targeting (top), or lncRAP2-targeting (bottom) DsiRNAs. (**d, e**) Coordinate changes in Igf2bp2 client RNA levels upon lncRAP2 or *Igf2bp2* depletion. RNA changes for Igf2bp2 targets upon *Igf2bp2* vs. lncRAP2 depletion in mature white adipocytes are shown in (d), highlighting differentially expressed (p <0.05) targets. Select ones are shown in (e). (**f**) Coordinate RNA and protein changes upon lncRAP2 depletion. Global protein vs. RNA changes upon lncRAP2 depletion in mature white adipocytes, highlighting Igf2bp2 targets with protein changes that are significant (p <0.05, red) or not (black). (**g, h**) lncRAP2-Igf2bp2 potentiate energy expenditure. Protein changes of Adipoq and Fabp4 upon lncRAP2 or *Igf2bp2* depletion in isoproterenol-stimulated or non-stimulated white adipocytes are shown in (g). Lipolysis assays measuring glycerol release in isoproterenol-stimulated or non-stimulated lncRAP2 or *Igf2bp2*-depleted white adipocytes are shown in (h). *p <0.05, **p <0.01, ***p <0.001 relative to negative control (t-test).

Adipogenesis induced or suppressed Igf2bp2 client RNAs are reciprocally regulated if lncRAP2 is depleted before differentiation^18^ (Supplementary Fig. 3d), consistent with common RNA targets. To study the lncRAP2-Igfb2p connection independent of effects on adipogenesis, we depleted them in mature adipocytes. Transducing DsiRNAs against lncRAP2 or *Igf2bp2* into differentiated white adipocytes deplete either by 70-80% without impacting the other’s levels (Fig. 3c). Unbiased RNA sequencing, however, reveals tightly correlated (r= 0.75) global gene expression changes upon lncRAP2 and *Igf2bp2* depletion (Fig. 3d). These include concordant reduction of mRNAs encoding Igf1/2 and thermogenic factors Ebf2 and PGC1α, while levels of adipocyte-specific non-target mRNAs are unaffected (Fig. 3c, e). Destabilization of mRNA targets is biased toward those encoding effectors of lipid/glucose synthesis and metabolism (Supplementary Fig. 3e), suggesting that the lncRAP2-Igf2bp2 complex normally potentiates energy expenditure. To test this, we assessed lipolysis in differentiated lncRAP2 and *Igf2bp2* – depleted adipocytes. In both cases, isoproterenol-induced lipolytic responses were reduced, and lncRAP2 depletion also lowered basal lipolysis (Fig. 3h). Supporting this, the gene signature of lysosomal acid lipase deficiency has the most significant association with that of lncRAP2/ *Igf2bp2* depletion (Supplementary Fig. 3f). Thus, lncRAP2-Igf2bp2 complexes support energy expenditure in mature adipocytes by binding and stabilizing mRNAs encoding many metabolic effectors.

Mechanistically, lncRAP2-Igf2bp2 complexes predominantly act to modulate the levels of client mRNAs, but do not directly affect their translation. To determine this, we assessed mRNA translation by quantitative mass spectrometry and ribosome footprint profiling. Proteins from 1,273 genes were quantifiable (by at least 2 unique peptides) in mature adipocytes using tandem mass tag labeling, including 138 Igf2bp2 targets. As with transcriptional changes, protein changes are tightly correlated (r= 0.73) upon lncRAP2/*Igf2bp2* depletion (Supplementary Fig. 4a), particularly for direct Igf2bp2 targets (r= 0.78), and proteins with roles in lipid/glucose metabolism and oxidation are selectively destabilized (Supplementary Fig. 4b), including Fabp4 and Adiponectin (Fig. 3g). Protein changes upon lncRAP2/*Igf2bp2* depletion are globally coupled with mRNA changes (Fig. 3f and Supplementary Fig. 4c), and this is true specifically for direct Igf2bp2 targets. Supporting this conclusion, we found no significant differences in the translation efficiency of Igf2bp2 mRNA targets vs. non-targets in primary adipocytes, whether down/up regulated at the protein level upon lncRAP2 or *Igf2bp2* depletion (Supplementary Fig. 4d). Thus, lncRAP2-Igf2bp2 complexes do not directly affect mRNA translation.

### lncRAP2 and Igf2bp2 are associated with obesity-linked diabetes risk

Given lncRAP2’s metabolic programming capacity, we sought validation from human genetics. A genetic variant (rs2209972:C) associated with increased body mass index, higher fasting insulin levels, and insulin resistance in women with polycystic ovary syndrome in a Chinese population^41^ maps within lncRAP2. While this variant has been associated to the neighboring insulin-degrading enzyme (IDE) gene^41^, we find that, in the Chinese or any other population surveyed, it is in strong linkage disequilibrium (r^2^≥ 0.8) with variants within lncRAP2 but no other loci within 500kb, including IDE (Fig. 4a and Supplemental Fig. 5a), confining its phenotypic associations to the lncRAP2 genetic locus.

**Fig. 4.**
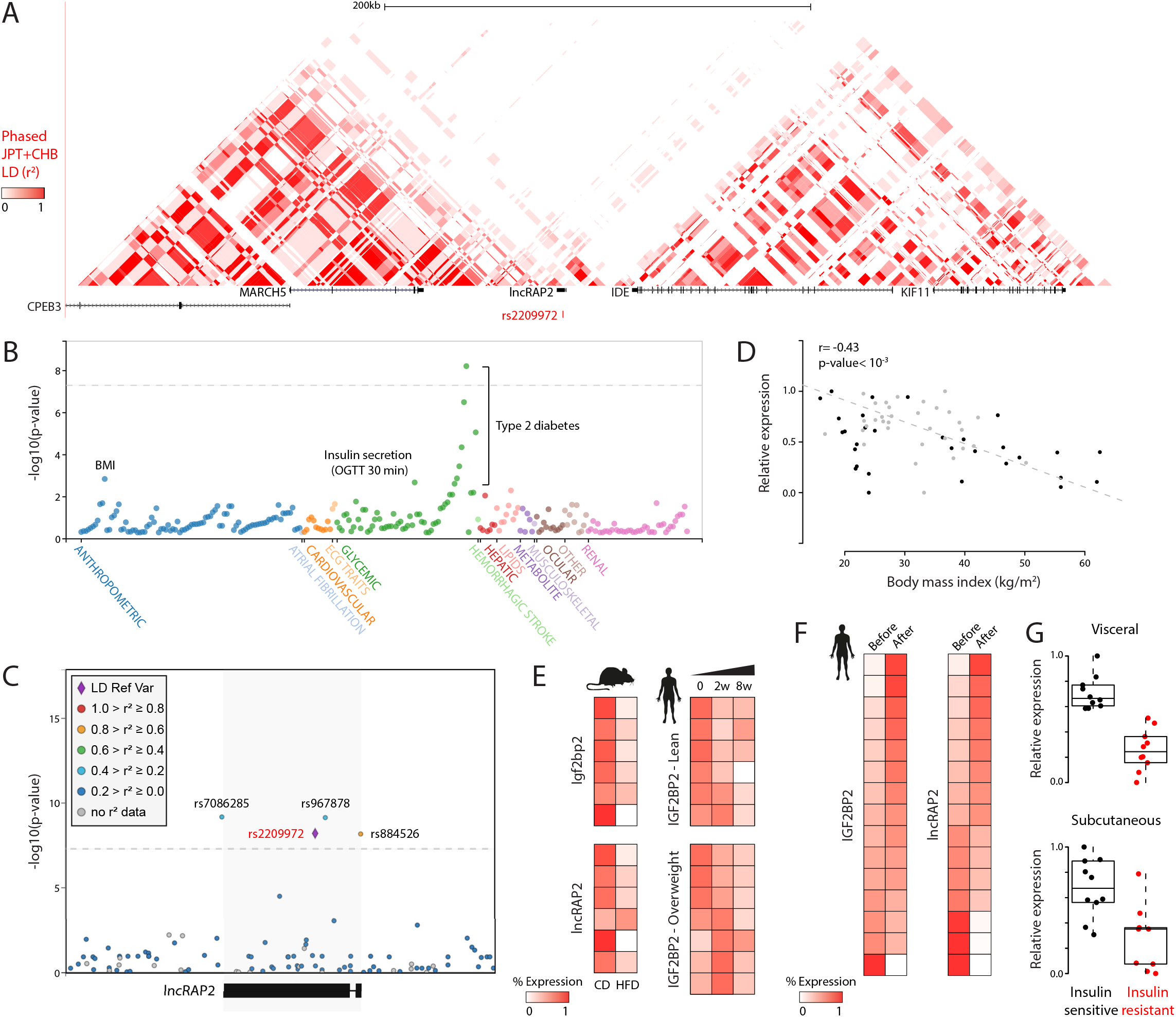
lncRAP2-Igf2bp2 genetic and expression variability are associated with obesity-linked diabetes risk. (**a**) Human lncRAP2 linkage disequilibrium block harbors rs2209972, a genetic variant linked to higher body mass, fasting insulin, and insulin resistance in Chinese females^41^. Track displays linkage disequilibrium (LD) for a population of Han Chinese in Beijing and Japanese in Tokyo (JPT+CHB)^72^ from phased genotypes of 90 unrelated individuals^73^. (**b**) lncRAP2 rs2209972 is specifically associated with body mass, insulin secretion, and type 2 diabetes. Phenome-wide association results between rs2209972 and 317 phenotypes across 16,278,030 individuals of various ancestries from the Type 2 Diabetes Knowledge Portal (http://www.type2diabetesgenetics.org/variantInfo/variantInfo/rs2209972). Only significant (p<0.05) associations are shown. (**c**) lncRAP2 locus-wide association with type 2 diabetes. Significance of association between genetic variants within and around the lncRAP2 locus (highlighted) and type 2 diabetes in 898,130 European-descent individuals^74^. Variants are colored based on 1000 Genomes All samples^75^ LD with rs2209972 (diamond). (**d, e**) Adipose lncRAP2 and *Igf2bp2* are progressively suppressed with obesity. Body mass index vs. *Igf2bp2* levels in visceral^76^ (black) or subcutaneous^77^ (gray) adipose tissue from 63 individuals shown in (d). Relative lncRAP2 and *Igf2bp2* expression in mice^78^ (visceral fat, left) or lean/overweight humans^79^ (subcutaneous fat, right) fed a high fat diet shown in (e). (**f**) Adipose lncRAP2 and *Igf2bp2* are restored with weight loss after bariatric surgery. Relative lncRAP2 and *Igf2bp2* expression in subcutaneous adipose tissue samples from 15 obese women before and 3 months after Roux-en-Y gastric bypass surgery^80^. (**g**) Adipose *Igf2bp2* is suppressed with insulin resistance. Relative *Igf2bp2* expression in both visceral (top) and subcutaneous (bottom) adipose tissue samples from 10 insulin-sensitive or 10 insulin-resistant body mass index-matched obese patients undergoing gastric bypass surgery^81^.

To probe rs2209972 associations with many different phenotypes without bias, we examined 99 genome-wide association studies of 234 traits. These included phenotypes relevant to metabolic, cardiovascular, hepatic, neural, renal, and musculoskeletal disorders, as well as anthropometric traits. Our analysis revealed specific enrichment of associations with body mass index, insulin secretion, and type 2 diabetes (Fig. 4b and Table 1). Of these associations, increased diabetes risk in individuals of European ancestry was the strongest (odds ratio= 1.04 per rs2209972:C allele, p< 10^−8^). Additional genetic variants mapping within lncRAP2 and its transcription start/end sites also represent risk alleles for type 2 diabetes (p< 10^−9^ to 10^−8^), thus evidencing lncRAP2-wide diabetes association (Fig. 4c and Table 1). Notably, Igf2bp2 is also specifically associated with the same metabolic traits (Supplementary Fig. 5b, c), in line with lncRAP2-Igf2bp2 complexes modulating a common genetic pathway.

Type 2 diabetes pathogenesis can involve obesity-linked adipose dysfunction^42,43^. To test lncRAP2-Igf2bp2’s contribution, we studied their regulation in obesity and diabetes progression. Adipose lncRAP2/*Igf2bp2* are progressively suppressed in mice/humans fed a high-fat diet (Fig. 4e), decrease with higher body mass index (Fig, 4d and Supplemental Fig. 5d) and are restored upon weight loss after bariatric surgery (Fig. 4f). Moreover, *Igf2bp2* is adipose-suppressed in leptin-deficient (ob/ob) and leptin receptor-deficient (db/db) mice (Supplemental Fig. 5e), which model diabetes onset and progression^44^. We further studied *Igf2bp2* in advanced diabetics, matched for body mass index, and found strong inhibition in visceral/subcutaneous adipose in insulin-resistant vs. sensitive patients (Fig. 4g). Thus, genetic and expression variation of lncRAP2-Igf2bp2 are associated with obesity-linked diabetes outcomes. We propose that this results from lncRAP2-Igf2bp2 complexes’ ability to program adipocyte metabolism by tuning mRNA levels of key energy expenditure genes.

## Discussion

Obesity has become pandemic^45,46^, increasing the worldwide prevalence of type 2 diabetes^47^. Anti-diabetic drugs targeting white fat have not been successful^48,49^, highlighting limited insight into how adipocytes develop and function. We and others have shown that lncRNAs contribute to adipocyte lineage specification and specialization^1,2^. Specific mechanisms have been elusive, however, due to poor conservation and scarcity of lncRNA-centric tools. In this study, we have made three important contributions to understanding how lncRAP2, conserved between mice and humans, regulates adipocyte function. First, genetic variants within lncRAP2 are specifically associated with diabetes, obesity, and metabolic traits. Second, lncRAP2 forms complexes with the RNA-binding protein Igf2bp2. And third, lncRAP2-Igf2bp2 complexes act to regulate mature adipocyte metabolism by stabilizing client mRNAs encoding energy metabolism effectors.

Our evidence shows that multiple genetic variants associated with propensity to type 2 diabetes map to a linkage disequilibrium block encompassing lncRAP2 but no other gene loci. Analysis of hundreds of phenotypes interrogated by genome-wide association studies reveals that the lncRAP2 alleles are specifically associated with body fat mass, insulin secretion, and type 2 diabetes, implicating lncRAP2 in the development of obesity and diabetes. Consistently, lncRAP2 is suppressed in the white fat of mice and humans fed a high fat diet, decreasing progressively with increasing obesity, whereas lncRAP2 levels are restored upon weight loss.

lncRAP2 is enriched in white over brown adipocytes and is critical for adipogenesis^18^. Although most lncRNAs appear to be nuclear-enriched^50^, lncRAP2 is mainly cytoplasmic, yet we find no evidence of productive translation by ribosome footprint profiling. Using RNA interactome analyses in cross-linked intact cells, we found that lncRAP2 does not directly bind other RNAs, but specifically interacts with metabolic enzymes and mRNA decay/translation modulators. Some of these appear to be transient/indirect interactions, which may be of importance in other tissues where lncRAP2 is detected. However, in mouse and human adipocytes, we verified strong, direct binding to endogenous Igf2bp2.

Igf2bp2 is of prime interest, as it harbors one of the first genetic variants associated with type 2 diabetes by genome-wide association studies^51–53^. This and other variants in linkage disequilibrium are associated with lower *Igf2bp2* in pancreatic islets^54^, and with impaired insulin responses^55^. Yet they are also associated with increased body fat^56^, with a stronger effect in type 2 diabetics^57^. Intriguingly, Igf2bp2-null mice rather resist diet-induced obesity and diabetes, though this is likely due to underdeveloped white and overactive brown fat^19^—a tissue that is much less prevalent in adult humans^58^. How Igf2bp2 impacts diet-induced obesity and diabetes risk, and its roles in mature white fat, have thus remained unclear. We find, in mature white adipocytes, Igf2bp2 globally binds and regulates mRNAs encoding key adipogenic regulators, explaining its requirement for *in vitro/in vivo* adipogenesis^19,20^, as well as energy metabolism effectors. The latter include Elovl factors, Fabp4, and Adiponectin, linking Igf2bp2 to lipid synthesis, transport, and metabolism. Indeed, we found hindered lipolytic responses upon *Igf2bp2* inhibition, signaling reduced energy expenditure capacity. Notably, metabolic mRNA targets in adipocytes are also bound by Igf2bp2 in human cells, and *Igf2bp2* suppression in visceral or subcutaneous fat distinguish insulin-resistant from-sensitive diabetics.

Remarkably, Igf2bp2 binds >55% of cellular lncRAP2 but not as a client RNA. Instead, lncRAP2 and Igf2bp2 interact to program white adipocyte development and metabolism. In support of this conclusion, depleting either lncRAP2 or Igf2bp2 in mature adipocytes results in tightly correlated transcriptome and proteome changes. mRNAs encoding energy metabolism effectors are selectively destabilized, in both cases limiting lipolytic activity. Moreover, Igf2bp2 client mRNA changes are coupled with protein but not translation efficiency changes in both cases, indicating that the lncRAP2-Igf2bp2 complex fine-tunes energy metabolism primarily by modulating client mRNA stability. Although inhibiting Igf2bp2 phenocopies the molecular and physiological effects of lncRAP2 inhibition, both interact with additional proteins, such that these effects must reflect perturbation of many functions in addition to those exerted in partnership with each other.

Our characterization of a lncRAP2-Igf2bp2 interaction as a posttranscriptional regulatory program in adipocyte metabolism echoes findings that the Airn and HIF1A-AS2 lncRNAs are functional Igf2bp2 cofactors in the developing brain^59^ and in cardiomyocytes^60^, respectively. Thus, Igf2bp2 appears to bind distinct lncRNAs to regulate distinct targets across diverse tissues. Given that lncRAP2 is adipose-specific and conserved in human, it represents an attractive target to selectively modulate Igf2bp2 activity within fat tissue to treat or prevent the progression of obesity-linked diabetes.

## Methods

### Mice

C57BL/6J mice were bred in house or purchased from Jackson Laboratories (stock # 000664). All mice were housed under a 12 hour light / dark cycle at constant temperature (20°C). All procedures were performed according to protocols approved by the Committee on Animal Care at the Massachusetts Institute of Technology.

### Isolation of primary cells and tissues

6 – 8 male 2 – 4 week old mice were sacrificed by CO2 asphyxiation and interscapular brown adipose tissue (BAT) and subcutaneous (inguinal) white adipose tissue (scWAT) was harvested into room temperature plain DMEM (Sigma, # 56499C). The fat pads were transferred into a well of a 6 well plate and minced with scissors for 5 minutes. Minced tissues were then transferred into a 50 ml conical tube with 3 ml Hank’s balanced salt solution (Gibco, # 14175-095) supplemented with 0.2 % collagenase A (Roche, # 10103578001) and 2 % BSA (Sigma-Aldrich, # A7906) using a 1 ml pipet tip with the tip cut off to allow aspiration of larger pieces. The tissues were incubated agitating (350rpm) and repeated vortexing every 5 minutes for 10 sec at 37°C for 30 min or 20 minutes for scWAT. Following collagenase digestion, 10 ml room temperature plain DMEM was added and cells were filtered through a 70 μm mesh filter (Corning, # 352350). Mature adipocytes and the stromal vascular fraction (SVF) were separated by centrifugation at 700 g for 5 min. The supernatant was removed and the SVF resuspended in 10 ml room temperature plain DMEM followed by additional filtering through a 30 μm mesh filter (Miltenyi Biotec # 130-041-407) and subsequent centrifugation at 700 g for 5 min. The SVF from subcutaneous white fat pads (scWAT) of 8 mice were then resuspended in 10 ml DMEM supplemented with 10 % heat inactivated new born calf serum (HI NCBS, LifeTechnologies / Gibco, # 26010-074) and plated on two 10 cm dishes (Corning, # 430293). The SVF from interscapular brown fat pads of 6 – 8 mice were then resuspended in 6 ml DMEM supplemented with 10 % HI NCBS and plated on 3 wells of a six well plate (Corning, #3506). After 4 and 24 h, the medium was replaced by fresh, pre-warmed DMEM / 10 % HI NBCS at 37°C and with 5 % CO_2_. Cells were grown to confluence and then passaged no more than two times before seeding the pre-adipocytes for differentiation.

### Cell culture

Pre-adipocytes derived from BAT were cultured to confluence and then subsequently overgrown for 4 – 6 additional days until growth arrested. The cells were then induced to differentiate by culturing them for two days in induction medium consisting of DMEM supplemented with 10 % fetal bovine serum (FBS, Sigma-Aldrich F2442) and 850 nM insulin (Sigma-Aldrich # I1882), 0.5 μM dexamethasone (Sigma-Aldrich # D4902), 250 μM 3-isobutyl-1-methylxanthine (IBMX, Sigma-Aldrich # I5879), 1 μM rosiglitazone (Cayman Chemical # 71742), 1 nM 3,3,5-triiodo-L-thyronine (T3, VWR # 100567-778) and 125 nM indomethacin (Sigma-Aldrich # I7378). Subsequently, the induction medium was replaced with DMEM supplemented with 10 % FBS and 160 nM insulin and 1 nM T3 for another two days. The cells were then cultured in DMEM 10 % FBS and 1 nM T3 until day 8 of differentiation, and the medium was replaced every other day. Pre-adipocytes from scWAT were cultured similar but the induction medium and following medium did not contain T3.

### Cell stimulation

Mature adipocyte cell layers were washed twice in plain pre-warmed DMEM and stimulated with 1 μM isoproterenol (Sigma-Aldrich I6504). After 6 h of stimulation, the cells were washed once with cold PBS and RNA was harvested using TRizol or QIAzol lysis reagent as described below. For immunoblotting, the cultures were harvested after stimulation with isoproterenol after washing with cold PBS on ice and adding 40 μl RIPA buffer per well of a 6 well plate or 100 μl RIPA buffer to a 10 cm dish.

### siRNA transfection

Was performed as described^22^. Briefly, on day 4 of differentiation, adipocytes were transfected with 5 nM siRNA (Sigma-Aldrich) using 5 μl/ml Lipofectamine RNAiMAX diluted in Opti-MEM I Reduced Serum Medium (Life Technologies). The cultures were analysed on day 6 of differentiation.

### Glycerol release

Following addition of fresh medium, cells were stimulated with isoproterenol. Cell culture medium was collected after 24h of stimulation and stored at −20°C. Glycerol release was measured using the Adipolysis Assay Kit (Cayman Chemical, 10009381) following the instructions of the manufacturer.

### Quantitative PCR

Total RNA was isolated from tissues or cells using TRizol or QIazol reagent (LifeTechnologies/Ambion) and a miRNAeasy kit (Qiagen). 300 ng were reverse transcribed using Superscript II reverse transcriptase (LifeTechnologies/Invitrogen) using random hexamers (LifeTechnologies/Invitrogen). The cDNA was diluted 1:10 and 2.5 μl for a 96 well plate or 1 μl for a 384 well plate were used for quantitative Real-time PCR. qPCR was carried out on an ABI7900HT Fast real-time PCR system (Applied Biosystems) and analysed using the delta delta Ct method normalized to 18S if not stated otherwise. Results are shown as pooled data from 3-4 independent experiments displaying the average and SEM.

### RNA sequencing analysis

Poly A+ RNA sequencing (TrueSeqStrandedPolyA) was performed on RNA samples using a HiSeq genome sequencer (Illumina). RNASeq paired-end reads from Illumina 1.5 encoding were aligned using TopHat (v 2.0.13)^61^ to the mouse genome (mm9; NCBI Build 37) with Ensembl annotation (release 67) in gtf format. mRNA sequencing was performed on three independent experiments. Differential analysis and counts per millions (cpm) were obtained using DESeq^62^ as described^31^.

### Immunoblotting

Lysates were centrifuged for 10 min at 13,000 g to remove debris, and NuPAGE sample buffer and reducing buffer (LifeTechnologies) were added after measuring and adjusting the samples for protein concentration (DC protein assay kit II, Bio-Rad). 2-20 μg protein per sample were separated for 2-4 h at 60 – 100 V using 8 % 26 well NuPAGE Bis-Tris Midi gels in MOPS or MES buffer (LifeTechnologies) and in Criterion cells (Bio-Rad) using respective adapters (LifeTechnologies). Protein was wet transferred to polyvinylidene fluoride (PVDF) membranes (Immobilon P, Millipore) in Criterion blotter cells (Bio-Rad) using 2 x NuPAGE Transfer buffer (LifeTechnologies) with 10 % methanol for 25 min at 1 A. After blocking the membrane in filtered (Nalgene # 595-4520) TBS-T (50 mM Trisbase, 150 mM NaCl, 0.1 % Tween 20 (Sigma-Aldrich # P1379)) with 3 % bovine serum albumin (BSA, Sigma-Aldrich # A7906) for 1 h, blots were incubated in primary antibody diluted in TBS-T 3 % BSA sealed in hybridization bags and gently shaking at 4°C overnight, then washed three times for 10 min in TBS-T, incubated in secondary antibody (Cell Signalling Technologies) diluted in TBS-T 3% BSA gently shaking for one hour at room temperature, and then washed again three times for 10 min in TBS-T. Antibody binding was visualized using ECL Plus Western Lightning reagent (Perkin Elmer NEL 102) and blots were exposed to film (Kodak BioMax MR Film, Carestream Health Inc # 8701302). Films were scanned without adjustments using an Epson scanner. All immunoblotting data shown were reproduced with almost identical results in at least one and typically two to three additional independent experiments.

### Oil-red O staining of brown adipocyte cultures

Cells were washed in PBS and fixed in 3.7% formaldehyde solution for 1 h, followed by staining with Oil Red O for 1 h. Oil Red O was prepared by diluting a stock solution (0.5 g of Oil Red O (Sigma) in 100 ml of isopropanol) with water (6:4) followed by filtration. After staining, plates were washed twice in water and photographed.

### 5’ and 3’ RACE

The 5’ and 3’ ends of lncRAP2 was determined using the FirstChoice RLM-Race Kit from Ambion following the manufacturers instructions. Primers were designed accordingly and can be found in Supplementary Data 3. Resulting gel bands were excised from the gel and purified using the Gel Extraction kit (Qiagen), cloned into a TopoTA vector (Thermo Fisher Scientific) and sequenced using the M13 fwd and rev primers supplied by the kit.

### Cell fractionation

To separate the nuclear from the cytoplasmic fraction differentiated 3T3-L1 adipocytes were harvested and 1 million cells were used to isolated the fractions using the PARIS kit (Life technologies) according to the manufacturers instructions. Separated fractions were then analyzed using Real-time PCR and gene specific primers.

### Single molecule FISH

Single molecule RNA FISH and fluorescence microscopy were described previously^26^. Briefly, antisense probes were designed to span the exons of lncRAP2 (Supplementary Data 3) and coupled to Cy5. Probes were hybridized at 2ng/μl final concentration. The maximum projection of FISH image z-stacks in the DAPI channel was merged with the z-slice of maximum contrast in the DIC channel, and the composite was used to identify cells. Images in the Cy5 channel were compared to those in the GFP control channel to detect diffraction-limited spots representing RNA transcripts using fixed pixel intensity thresholds. For image presentation, enhanced contrast in the DAPI channel was used to emphasize nuclear counterstaining boundaries.

### RNA Coimmunoprecipitation (RIP)

RNA coimmunprecipitation was done as described^63^. Mature adipocytes were harvested using trypsin. 1×10^7^ re-suspended in 2ml 1X PBS, then lysed in nuclear isolation buffer (2ml nuclear isolation buffer + 6ml water, premixed) for 20 minutes. Nuclei were pelleted by centrifugation at 2,500 g for 15 minutes. Supernatant was discarded and nuclei were re-supended in 1ml RIP buffer containing the HALT protease and phosphatase inhibitor (Thermo scientific) and split into two fractions and mechanically sheared using a dounce homogenizer with 20 strokes. Nuclear membrane and debris were pelleted by centrifugation at 13,000 rpm for 10 minutes at 4°C. The supernatant was pre-cleared by adding 30μl slurry of protein A/G beads (Santa Cruz, sc-2003) and incubation for 2 hours at 4°C on a rotator. Beads were removed by centrifugation at 2500 g for 1 minute and 10% of the supernatant was removed to a new tube (10% input) and the rest was incubated with antibodies to Suz12 (6μg; abcam, ab12073), IgG (10μg; abcam, ab37415), Ezh2 (8μg; abcam, ab3748), hnRNPU (8μg; abcam, ab20666), CoREST (8μg, santa cruz, sc23449) or no antibody for 3h at 4°C on a rotator. Then 60μl slurry of protein A/G beads were added for 2h at 4°C on a rotator. Beads were pelleted by centrifugation at 2500 rpm for 30s and washed 3 times in 500μl RIP for 10 minutes each followed by one wash with 1X PBS. For the isolation of RNA, the beads were re-suspended in TRIzol after the last wash step and isolated according to the manufacturers instructions. The RNA pellet was re-suspended in 10μl dH_2_O and was directly used for reverse transcription using random hexamers and SuperScript II (Invitrogen). Analysis was done by real time PCR and sequencing.

### RNA interactome analysis followed by sequencing (RIA-seq)

RNA FISH probes were biotinylated using Biotin-XX, SSE (Thermo Fisher). As controls, we also used probes against H19 (Supplementary Data 3) and Bloodlinc^31^. RIA-seq was conducted as described^11,28^. Briefly, 40 million day 6 differentiated adipocytes (3T3-L1) cells were harvested and crosslinked with 1% Glutaraldehyde for 10min at room temperature on a shaker, and the reaction was quenched with 0.125M glycine for 5min. Cells were collected by spinning at 2000g for 5min, resuspended in 4ml ice-cold PBS, and aliquoted to 4 collection tubes followed by resuspension in Lysis Buffer supplemented with Halt Protease and phosphatase inhibitor cocktail (Thermo Fisher, 100X) and with SUPERaseIn RNase Inhibitor (Life Technologies). Lysates were immediately sonicated in a Bioruptor (Diagenode) at 4°C using highest settings with 30 seconds ON and 45 seconds OFF for 3:45 hours. The sonicated samples were spun at 16100g for 10min at 4°C, and the supernatant was then transferred to a new tube, with 10% taken as input RNA control (in Trizol). Hybridization Buffer and 100pmol of the probes were added to the supernatant and incubated at 37°C for 4h in a rotator. 100μl of pre-washed Dynabeads MyOne Streptavidin C1 magnetic beads (Life Technologies) were then added and incubated for 30 minutes at 37°C in a rotator, captured by magnets (Invitrogen), and washed with Wash Buffer for 5 minutes at 37°C for a total of five times. After the last wash, beads were resuspended in 1ml Wash Buffer and then lysed using TRIzol.

### RIA-seq analysis

Raw reads (40bp unpaired and non-strand-specific) were mapped to mm9 using bowtie2^64^. Peak calling was performed on uniquely-mapped reads relative to input control using MACS (Zhang et al. 2008) with default parameters and “--bw=300 --mfold 10 30”. High-confidence peaks were identified based on several criteria. First, peak coverage was quantified using BEDTools (Quinlan and Hall 2010), and only peaks with an average per-base coverage greater than 2 reads were considered. Then peaks whose read density enrichment was at least 10-fold greater than the background model were considered enriched. Finally, peaks in the experimental sample overlapping peaks in the input control were excluded from further analysis.

### Comprehensive identification of RNA-binding proteins by mass spectrometry (ChIRP-MS)

ChIRP-MS was performed as described^14^ with the biotinylated probes used for RIA-seq. Since the first capture using even and odd probes didn’t result in pull-downs, we used all 48 exon probes for pull-down and intron, Bloodlinc, H19 probes as well as no probe control and RNAse A treated samples as control. Briefly, 100 million day 6 differentiated adipocytes from 3T3-L1 cells were harvested and washed twice in PBS, crosslinked in 3% formaldehyde for 30min, then quenched with 0.125M glycine for 5min, and collected by spinning at 2000g for 5 min. Cells were lysed and sonicated for 2:30 hours. Lysates were pre-cleared by incubating with 50μl washed magnetic beads at 37°C for 30min and retrieving them with magnets before proceeding to hybridization with probes overnight at 37°C in a rotator. The following day, beads were added and incubated for 30 minutes at 37°C in a rotator, then washed five times with Wash Buffer but not eluted as described in ^14^; instead, beads were boiled for 10min at 95°C in Laemmli Buffer, then beads and buffer were separated on a NuPAGE 4-12% Bis-Tris gel, followed by silver staining to identify differential bands. The whole gel lane was then excised, trypsinized, reduced, alkylated, and trypsinized at 37°C overnight. The resulting peptides were extracted, concentrated, and HPLC-purified. Effluents were analyzed using an Orbitrap Elite (ThermoFisher) mass spectrometer in nanospray configuration operated in data-dependent acquisition mode, where the 10 most abundant peptides detected using full scan mode with a resolution of 240,000 were subjected to daughter ion fragmentation in the linear ion trap. A running list of parent ions was tabulated to an exclusion list to increase the number of peptides analyzed throughout the chromatographic run.

### ChIRP-MS analysis

Mass spectrometry fragmentation spectra were correlated against a custom peptide database, formed by downloading all RefSeq species-specific (mm9) entries, and against a database of common contaminants (keratin, trypsin, etc.) using Mascot (Matrix Science) 2.5.1 and the Sequest algorithm (Thermo). The resulting Mascot search results were uploaded into Scaffold (Proteome Software), and a minimum of two peptides and peptide threshold of 95% and protein threshold of 99% were used for identification of peptides and positive protein identifications. Proteins enriched in ChIRP-MS experiments were identified based on several criteria. First, only proteins identified by 2 or more unique peptides in the differentiated sample were considered. Then proteins whose total peptide count was at least 10-fold greater in the exon probes versus intron probes were considered enriched. Finally, proteins introduced during sample preparation and purification (e.g. streptavidin, albumin) were excluded from further analysis. To distinguish lncRAP2-specific interactors from proteins that generally bind RNA during its processing, we compared the proteins enriched by lncRAP2 with those enriched by Dleu2^31^, Neat1 and Malat1^29^, and U1, U2, and Xist^14^, which were also filtered for identification by 2 or more unique peptides, for enrichment relative to their control sample greater than 3-fold, and for known contaminants. Proteins enriched by each RNA were ranked according to their total peptide counts, and rankings were compared.

### Silver staining

After proteins were separated on an Acryl-amid gel (see western blot), the gel was fixed in 40% EtOH with 10% HAc for 1 hour then washed 2×20 minutes in 30% EtOH and once for 20 minutes in water. The gel was sensitized for 1 minute in 0.02% Na2S2O3 following 3×20 seconds wash in water and 20 minutes incubation in cold (4°C) 0.1% AgNO3 solution. The gel was washed 3×20 seconds in water and developed in a new chamber in 3% Na2CO3 with 0.05% formaldehyde. As soon as bands became visible the reaction was stopped by a short wash in water and then incubation in 5% HAc. All lanes were cut out and processed (see ChIRP-MS analysis).

### Gene set and pathway enrichment analysis

Gene lists were analyzed for enrichment of genes grouped by biological process ontology or by curated annotations from the Molecular signatures database with GSEA^65^ using default parameters and “-metric log2_Ratio_of_Classes”.

### Motif enrichment analysis

Genomic sequences from regions of interest (e.g. UTRs) were searched for matches to a database of TF recognition sites^66^ for TFs expressed in the relevant cell type using FIMO ^67^ as described in^68^ with minor modifications: a Markov model of sequence nucleotide composition was used as the background model for motif matching (to normalize for biased distribution of individual letters in the examined sequences), and motifs with an odds ratio>2 and q-value<0.05 (Fisher’s exact test) relative to 10 randomly-shuffled controls were considered significantly enriched.

### Additional bioinformatics methods

All sequencing reads were quality-checked with FastQC (http://www.bioinformatics.babraham.ac.uk/projects/fastqc/). Genome-wide read density maps were generated by MACS2 using the “--bdg” option, normalized by RSeQC^69^ using the “normalize_bigwig.py” function, and visualized using BEDTools and the UCSC genome browser. Signal coverage and signal change surrounding regions of interest (e.g. DMRs, enhancer sites) were visualized using the ngs.plot R package ^70^. Data heatmaps were generated using the heatmap.2 function of the gplots R package (http://CRAN.R-project.org/package=gplots).

### Statistical methods

No statistical methods were used to predetermine sample size or remove outliers. The statistical difference between two sets of paired count data (e.g. motif matches in test vs. randomly-shuffled sequences) was assessed by a Fisher’s exact test using the fisher.test R implementation with default parameters. For unpaired data, a Shapiro-Wilk normality test was first performed using the shapiro.test R implementation with default parameters; for normally distributed data we then used a two-sided t-test (t.test R implementation with default parameters) to assess confidence on the measured difference of their mean values. For unpaired data that don’t follow a normal distribution, we used a non-parametric Wilcoxon rank sum test to determine if they belong to the same distribution. Variance was represented as mean ±SEM of n=3 replicates unless otherwise specified.

## Acknowledgements

We thank the imaging, proteomics, bioinformatics, and genome core facilities at the Whitehead Institute for technical support. We also thank Dr. Alexander Bartelt and Dr. Wenqian Hu for critical discussions. J.R.A.-D. is a Howard Hughes Medical Institute Fellow of the Life Sciences Research Foundation. This research was supported by fellowship Kn1106/1-1 from the Deutsche Forschungsgemeinschaft to M.K. and National Institute of Health grants DK068348 and DK047618 to H.F.L

## Author contributions

J.R.A.-D., M.K., and S.W. performed experiments; J.R.A.-D., M.K., and H.F.L. designed the research, interpreted results, and wrote the manuscript.

---

**Supplementary Fig. 1 lncRAP2 is a conserved cytoplasmic RNA required for adipogenesis.**

(**a**) lncRAP2 structure, expression, and regulation in brown adipocytes. Tracks show signal from sequencing studies of primary brown adipocytes from mouse (a) and human^71^ (b).

(**b**) Rapid amplification of cDNA ends (RACE) delineates the lncRAP2 transcript.

**Supplementary Fig. 2 lncRAP2 forms a complex with proteins that regulate mRNA stability and translation.**

(**a**) Specific and reproducible enrichment of spliced lncRAP2 by antisense purification. Tracks show signal from RNA interactome analysis by sequencing (RIA-seq) studies of differentiated white adipocytes. Locations of lncRAP2 exon- and intron-targeting antisense probe pools are shown below.

(**b**) lncRAP2 does not directly bind other RNAs. A minority of RIA-seq peaks are common to both lncRAP2 exon-targeting probe pools, and show poor enrichment or concordance between the probe pools, other than the peaks from capturing lncRAP2 itself.

(**c**) Specific enrichment of protein analytes in differentiated 3T3-L1 adipocytes recovered by lncRAP2 antisense purification. Silver stain of captured proteins before mass spectrometry analysis is shown.

(**d**) Specific native immunoprecipitation of Igf2bp2 (highlighted) in primary mouse white adipocytes. Western blot of captured proteins from an aliquot after pulldown with an Igf2bp2-specific antibody.

(**e**) *Igf2bp2* expression in selected tissues showing enrichment in adipose tissue. Relative expression in different organs and tissues from mouse^4^ (top) and human^82^ (bottom).

(**f**) *Igf2bp2* expression during *in vitro* differentiation of white or brown adipocytes from mouse (left) and human^71^ (right).

(**g**) Igf2bp2 protein levels during *in vitro* differentiation of mouse white or brown adipocytes. Western blots for the indicated proteins are shown.

**Supplementary Fig. 3 lncRAP2-Igf2bp2 target transcripts of metabolic effectors to potentiate energy expenditure.**

(**a**) Igf2bp2 selectively binds mRNAs encoding adipogenic regulators and effectors. Enrichment of RNAs in native Igf2bp2 or control immunoprecipitates (n=3 replicates each) in mouse white adipocytes.

(**b**) Igf2bp2 targets enrich for the known Igf2bp2 binding motif predominantly within 3’UTRs of target mRNAs. Analysis of short motifs and their relative enrichment within 5’UTR, coding sequence, and 3’UTR regions of transcripts bound by Igf2bp2 in mouse white adipocytes. The top DREME^83^ motif identified (bottom) and the known Igf2bp2 binding motif^38^ (top) are compared.

(**c**) Most Igf2bp2 targets in mouse white adipocytes also copurify with Igf2bp2 in human cells. Overlap between targets identified by RNA immunoprecipitation sequencing (RIP-seq) in murine white adipocytes, by enhanced crosslinking and immunoprecipitation (eCLIP) in human embryonic stem (H9)^39^ and erythroleukemia (K562) cells^40^, and by photoactivatable ribonucleoside-enhanced crosslinking and immunoprecipitation (PAR-CLIP) in human embryonic kidney (HEK 293) cells^33^. Common adipogenic regulator and effector targets are highlighted.

(**d**) Induction and suppression of Igf2bp2 RNA clients with adipogenesis is reversed in lncRAP2-depleted cells. Client RNA changes in day 8 differentiated untreated vs. lncRAP2-depleted murine white adipocytes. Genes significantly (p <0.05) upregulated (red) or downregulated (green) upon lncRAP2 depletion are highlighted.

(**e**) lncRAP2/*Igf2bp2* depletion cause coordinate destabilization of transcripts encoding mediators of energy metabolism. Gene set enrichment analysis highlighting significant (p <0.05) biological processes upon *Igf2bp2* vs. lncRAP2 depletion.

(**f**) Gene set enrichment analysis reveals significant overlap between the lncRAP2 and *Igf2bp2* depletion gene signatures and that of lysosomal acid lipase deficiency. NES, normalized enrichment score.

**Supplementary Fig. 4 lncRAP2-Igf2bp2 predominantly tune the levels of target mRNAs.**

(**a**) Coupled protein changes upon lncRAP2 or *Igf2bp2* depletion. Global protein changes upon *Igf2bp2* vs. lncRAP2 depletion in mature white adipocytes, highlighting Igf2bp2 targets with protein changes that are significant (p <0.05, red) or not (black).

(**b**) lncRAP2 and *Igf2bp2* depletion cause coordinate reduction in levels of key energy metabolism effector proteins. Gene set enrichment analysis highlighting significant (p <0.05) biological processes upon *Igf2bp2* vs. lncRAP2 depletion in mature white adipocytes.

(**c**) Coordinate RNA and protein changes upon *Igf2bp2* depletion. Global protein vs. RNA changes upon *Igf2bp2* depletion in mature white adipocytes, highlighting Igf2bp2 targets with protein changes that are significant after Igfbp2 depletion (p <0.05, red) or not (black).

(**d**) lncRAP2 or *Igf2bp2* depletion do not alter Igf2bp2 translation efficiency of client mRNAs. Translation efficiency, measured by the enrichment of ribosome footprint profiling over RNA-seq reads in visceral adipose tissue^24^, for Igf2bp2 targets and non-targets upregulated (top) or downregulated (bottom) at the protein level upon lncRAP2/*Igf2bp2* depletion in mature white adipocytes.

**Supplementary Fig. 5 lncRAP2-Igf2bp2 genetic and expression variability are associated with obesity-linked diabetes risk.**

(**a**) Human rs2209972 is in strong linkage disequilibrium with genetic variants within lncRAP2 but no other loci within 500kb. Linkage disequilibrium with rs2209972 (diamond) based on 1000 Genomes^75^ data for the indicated populations.

(**b**) Regional association with type 2 diabetes at the IGF2BP2 locus. Significance of association between genetic variants surrounding the IGF2BP2 transcription start site and type 2 diabetes in 898,130 European-descent individuals^74^. Variants are colored based on 1000 Genomes All samples^75^ linkage disequilibrium with rs7615045 (diamond).

(**c**) IGF2BP2 rs7615045 is specifically associated with body mass, glycated hemoglobin (HbA1c), and type 2 diabetes. Phenome-wide association results between rs7615045 and 317 phenotypes across 16,278,030 individuals of various ancestries from the Type 2 Diabetes Knowledge Portal (http://www.type2diabetesgenetics.org/variantInfo/variantInfo/rs7615045). Only significant (p<0.05) associations are shown.

(**d**) Adipose lncRAP2 is progressively reduced with obesity. lncRAP2 levels in adipose tissue samples from patients with different body mass index (BMI)

(**e**) *Igf2bp2* is suppressed during diabetes onset and progression. *Igf2bp2* levels in visceral adipose tissue from leptin-deficient (ob/ob) and leptin receptor-deficient (db/db) 9-10 week-old mice^84^.

**Supplementary Data 1.** High-confidence lncRAP2-associated proteins and their peptide counts.

**Supplementary Data 2.** Enriched Igf2bp2-associated RNAs and their enrichment values.

**Supplementary Data 3.** Oligos and antibodies used in this study.

**Supplementary Data 4.** Datasets used in this study.

