## Supplemental Figures for "An adipocyte-specific lncRAP2 – Igf2bp2 complex enhances adipogenesis and energy expenditure by stabilizing target mRNAs"

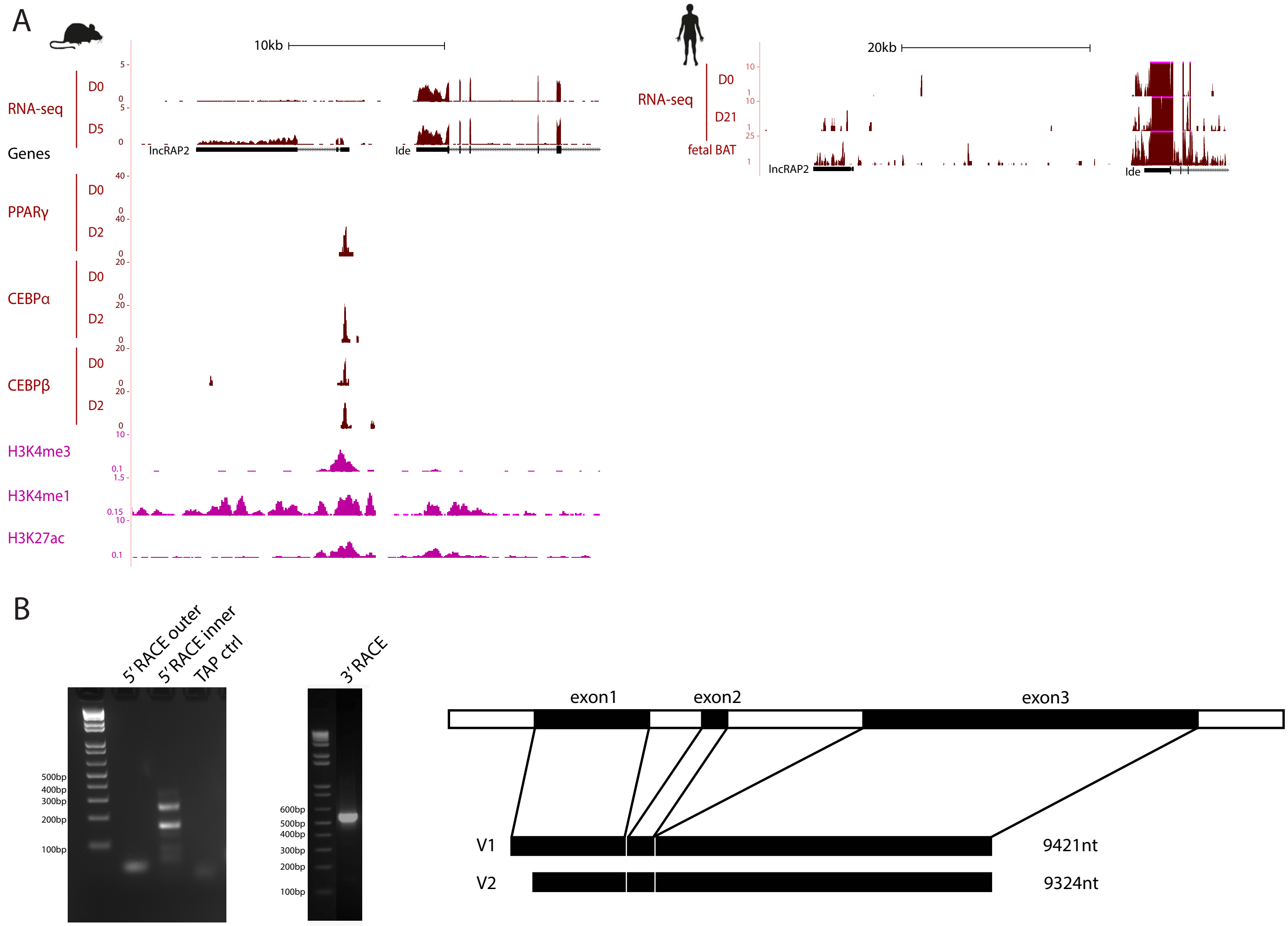
A
10kb
RNA-seq
D0
D5
Genes
lncRAP2
Ide
PPARγ
D0
D2
CEBPα
D0
D2
CEBPβ
D0
D2
H3K4me3
H3K4me1
H3K27ac
20kb
RNA-seq
D0
D21
fetal BAT
lncRAP2
Ide
B
5' RACE outer
5' RACE inner
TAP ctrl
500bp
400bp
300bp
200bp
100bp
3' RACE
600bp
500bp
400bp
300bp
200bp
100bp
exon1
exon2
exon3
V1
9421nt
V2
9324nt

**Supplementary Fig. 1 lncRAP2 is a conserved cytoplasmic RNA required for adipogenesis.**

(**a**) lncRAP2 structure, expression, and regulation in brown adipocytes. Tracks show signal from sequencing studies of primary brown adipocytes from mouse (a) and human[71] (b).

(**b**) Rapid amplification of cDNA ends (RACE) delineates the lncRAP2 transcript.

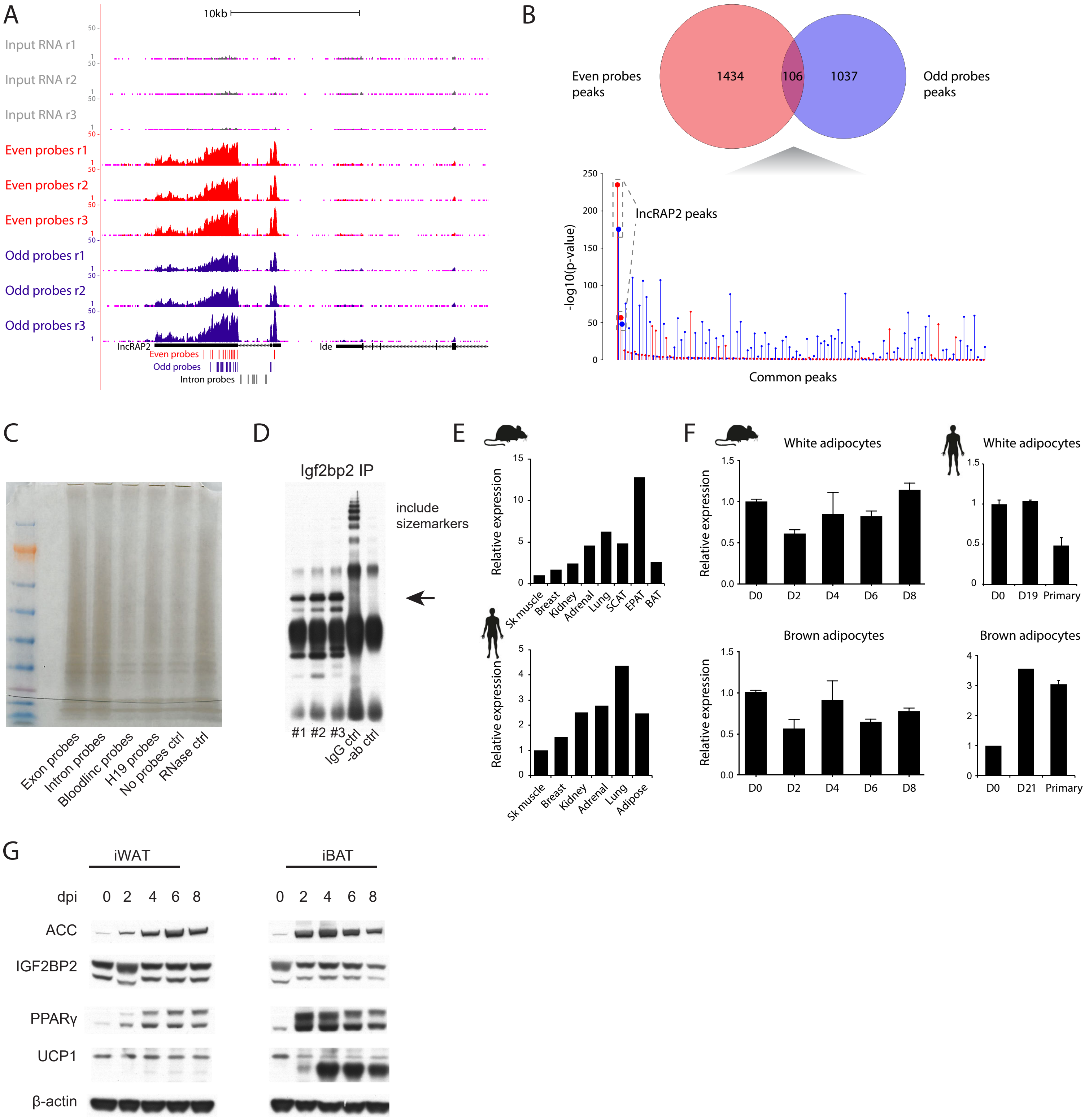
A
10kb
Input RNA r1
Input RNA r2
Input RNA r3
Even probes r1
Even probes r2
Even probes r3
Odd probes r1
Odd probes r2
Odd probes r3
lncRAP2
Ide
Even probes
Odd probes
Intron probes
B
Even probes peaks
1434
106
1037
Odd probes peaks
-log10(p-value)
lncRAP2 peaks
Common peaks
C
Exon probes
Intron probes
Bloodlinc probes
H19 probes
No probes ctrl
RNase ctrl
D
Igf2bp2 IP
include sizemarkers
#1 #2 #3
IgG ctrl
-ab ctrl
E
Relative expression
Sk muscle
Breast
Kidney
Adrenal
Lung
SCAT
EPAT
BAT
Adipose
F
White adipocytes
Brown adipocytes
D0
D2
D4
D6
D8
D19
D21
Primary
G
iWAT
iBAT
dpi
0 2 4 6 8
ACC
IGF2BP2
PPARγ
UCP1
β-actin

**Supplementary Fig. 2 lncRAP2 forms a complex with mRNA stability and translation regulators.**

(**a**) Specific and reproducible enrichment of spliced lncRAP2 by antisense purification. Tracks show signal from RNA interactome analysis by sequencing (RIA-seq) studies of differentiated white adipocytes. Locations of lncRAP2 exon- and intron-targeting antisense probe pools are shown below.

(**b**) lncRAP2 does not directly bind other RNAs. A minority of RIA-seq peaks are common to both lncRAP2 exon-targeting probe pools, and show poor enrichment or concordance between the probe pools, other than the peaks from capturing lncRAP2 itself.

(**c**) Specific enrichment of protein analytes by lncRAP2 antisense purification in differentiated 3T3-L1 adipocytes. Silver stain of captured proteins before mass spectrometry analysis is shown.

(**d**) Selective purification of native Igf2bp2 (highlighted) in mouse white adipocytes. Western blot of captured proteins after pulldown with an Igf2bp2-pecific antibody before analysis of associated RNAs is shown.

(**e**) *Igf2bp2* adipose tissue enrichment. Relative expression in different organs and tissues from mouse[18] (top) and human[70] (bottom).

(**f**) *Igf2bp2* expression during *in vitro* differentiation of white or brown adipocytes from mouse (left) and human[71] (right).

(**g**) Igf2bp2 protein levels during *in vitro* differentiation of mouse white or brown adipocytes. Western blot for the indicated proteins is shown.

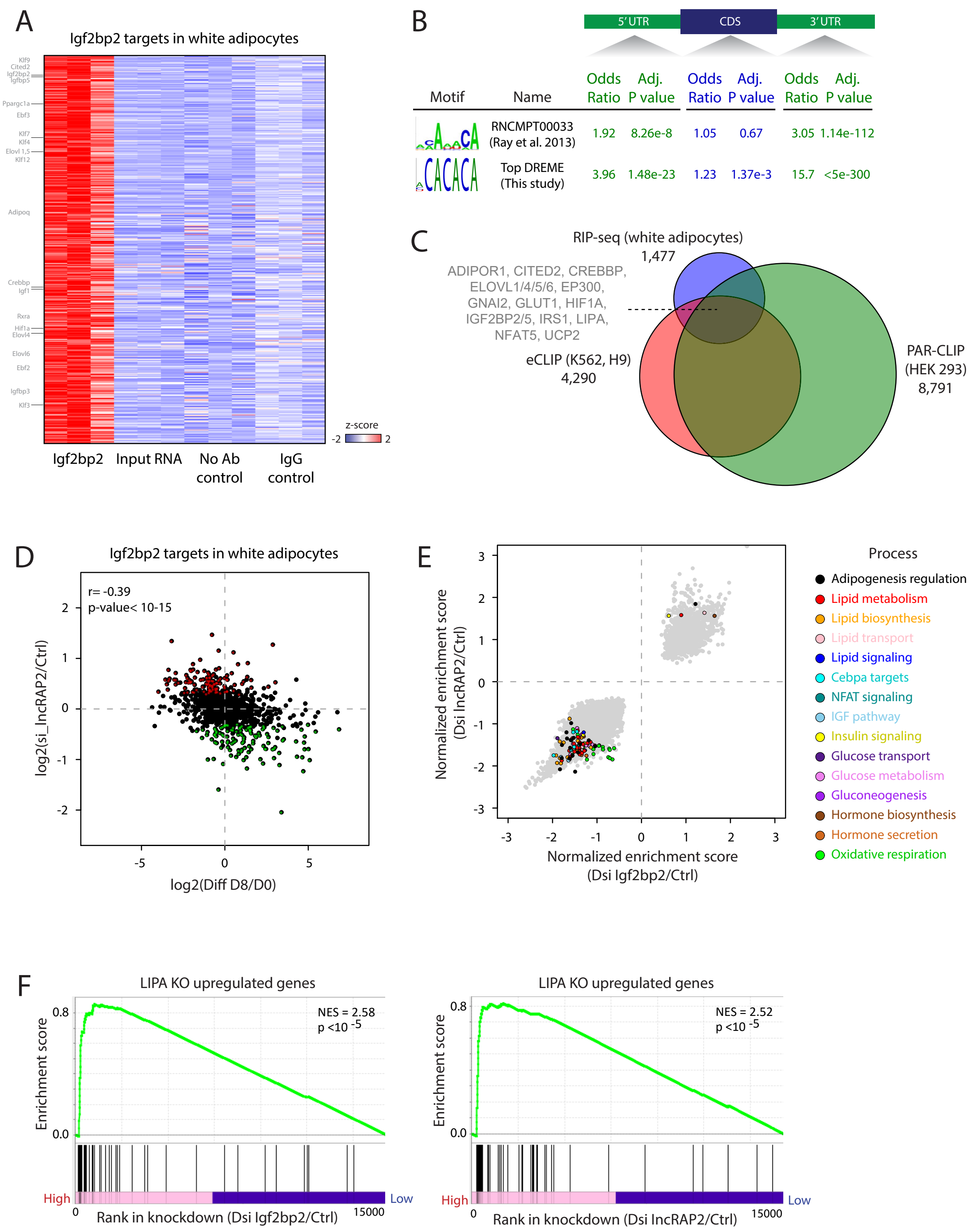
A
Igf2bp2 targets in white adipocytes
z-score
-2
2
Igf2bp2
Input RNA
No Ab control
IgG control
B
5′ UTR
CDS
3′ UTR
Motif
Name
Odds Ratio
Adj. P value
Odds Ratio
Adj. P value
Odds Ratio
Adj. P value
RNCMPT00033 (Ray et al. 2013) 1.92 8.26e-8 1.05 0.67 3.05 1.14e-112
Top DREME (This study) 3.96 1.48e-23 1.23 1.37e-3 15.7 <5e-300
C
RIP-seq (white adipocytes)
1,477
ADIPOR1, CITED2, CREBBP, ELOVL1/4/5/6, EP300, GNAI2, GLUT1, HIF1A, IGF2BP2/5, IRS1, LIPA, NFAT5, UCP2
eCLIP (K562, H9)
4,290
PAR-CLIP (HEK 293)
8,791
D
Igf2bp2 targets in white adipocytes
r= -0.39
p-value< 10-15
log2(si_lncRAP2/Ctrl)
log2(Diff D8/D0)
E
Normalized enrichment score (Dsi lncRAP2/Ctrl)
Normalized enrichment score (Dsi Igf2bp2/Ctrl)
Process
Adipogenesis regulation
Lipid metabolism
Lipid biosynthesis
Lipid transport
Lipid signaling
Cebpa targets
NFAT signaling
IGF pathway
Insulin signaling
Glucose transport
Glucose metabolism
Gluconeogenesis
Hormone biosynthesis
Hormone secretion
Oxidative respiration
F
LIPA KO upregulated genes
NES = 2.58
p <10 -5
Enrichment score
High
Low
Rank in knockdown (Dsi Igf2bp2/Ctrl)
15000
LIPA KO upregulated genes
NES = 2.52
p <10 -5
Enrichment score
High
Low
Rank in knockdown (Dsi lncRAP2/Ctrl)
15000

**Supplementary Fig. 3 lncRAP2-Igf2bp2 target transcripts of metabolic effectors to potentiate energy expenditure.**

(**a**) Igf2bp2 selectively binds mRNAs encoding adipogenic regulators and effectors. Enrichment of RNAs in native Igf2bp2 or control immunoprecipitates (n=3 replicates each) in mouse white adipocytes.

(**b**) Igf2bp2 targets enrich for the known Igf2bp2 binding motif predominantly within 3'UTRs of target mRNAs. Analysis of short motifs and their relative enrichment within 5'UTR, coding sequence, and 3'UTR regions of transcripts bound by Igf2bp2 in mouse white adipocytes. The top DREME[80] motif identified (bottom) and the known Igf2bp2 binding motif[35] (top) are compared.

(**c**) Most Igf2bp2 targets in mouse white adipocytes also copurify with Igf2bp2 in human cells. Overlap between targets identified by RNA immunoprecipitation sequencing (RIP-seq) in murine white adipocytes, by enhanced crosslinking and immunoprecipitation (eCLIP) in human embryonic stem (H9)[36] and erythroleukemia (K562) cells[37], and by photoactivatable ribonucleoside-enhanced crosslinking and immunoprecipitation (PAR-CLIP) in human embryonic kidney (HEK 293) cells[30]. Common adipogenic regulator and effector targets are highlighted.

(**d**) Induction and suppression of Igf2bp2 RNA clients with adipogenesis is reversed in lncRAP2-depleted cells. Client RNA changes in day 8 differentiated untreated vs. lncRAP2-depleted murine white adipocytes. Genes significantly ($p < 0.05$) upregulated (red) or downregulated (green) upon lncRAP2 depletion are highlighted.

(**e**) lncRAP2/*Igf2bp2* depletion cause coordinate destabilization of transcripts encoding mediators of energy metabolism. Gene set enrichment analysis highlighting significant ($p < 0.05$) biological processes upon *Igf2bp2* vs. lncRAP2 depletion.

(**f**) Gene set enrichment analysis reveals significant overlap between the lncRAP2 and *Igf2bp2* depletion gene signatures and that of lysosomal acid lipase deficiency. NES, normalized enrichment score.

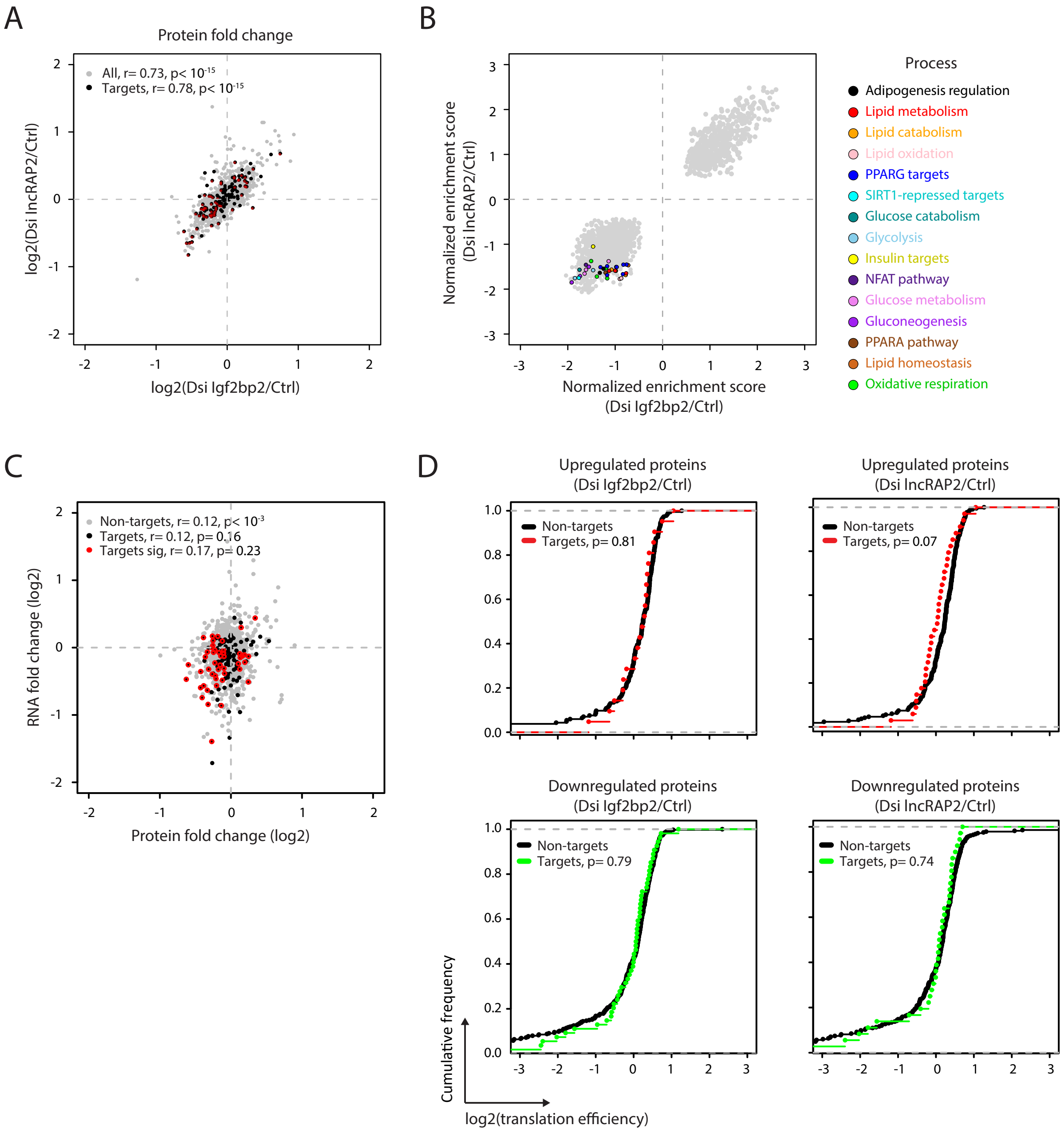
A
Protein fold change
All, r= 0.73, p< 10-15
Targets, r= 0.78, p< 10-15
log2(Dsi lncRAP2/Ctrl)
log2(Dsi Igf2bp2/Ctrl)
B
Normalized enrichment score
(Dsi lncRAP2/Ctrl)
Normalized enrichment score
(Dsi Igf2bp2/Ctrl)
Process
Adipogenesis regulation
Lipid metabolism
Lipid catabolism
Lipid oxidation
PPARG targets
SIRT1-repressed targets
Glucose catabolism
Glycolysis
Insulin targets
NFAT pathway
Glucose metabolism
Gluconeogenesis
PPARA pathway
Lipid homeostasis
Oxidative respiration
C
Non-targets, r= 0.12, p< 10-3
Targets, r= 0.12, p= 0.16
Targets sig, r= 0.17, p= 0.23
RNA fold change (log2)
Protein fold change (log2)
D
Upregulated proteins
(Dsi Igf2bp2/Ctrl)
Non-targets
Targets, p= 0.81
Upregulated proteins
(Dsi lncRAP2/Ctrl)
Non-targets
Targets, p= 0.07
Downregulated proteins
(Dsi Igf2bp2/Ctrl)
Non-targets
Targets, p= 0.79
Downregulated proteins
(Dsi lncRAP2/Ctrl)
Non-targets
Targets, p= 0.74
Cumulative frequency
log2(translation efficiency)

**Supplementary Fig. 4 lncRAP2-Igf2bp2 predominantly tune target mRNA levels.**

(**a**) Coupled protein changes upon lncRAP2/*Igf2bp2* depletion. Global protein changes upon *Igf2bp2* vs. lncRAP2 depletion in mature white adipocytes, highlighting Igf2bp2 targets with protein changes that are significant ($p < 0.05$, red) or not (black).

(**b**) lncRAP2/*Igf2bp2* depletion cause coordinate destabilization of energy metabolism effector proteins. Gene set enrichment analysis highlighting significant ($p < 0.05$) biological processes upon *Igf2bp2* vs. lncRAP2 depletion in mature white adipocytes.

(**c**) Coordinate RNA and protein changes upon *Igf2bp2* depletion. Global protein vs. RNA changes upon *Igf2bp2* depletion in mature white adipocytes, highlighting Igf2bp2 targets with protein changes that are significant ($p < 0.05$, red) or not (black).

(**d**) lncRAP2/*Igf2bp2* depletion do not alter Igf2bp2 client translation efficiency. Translation efficiency, measured by the enrichment of ribosome footprint profiling over RNA-seq reads in visceral adipose tissue[17], for Igf2bp2 targets and non-targets upregulated (top) or downregulated (bottom) at the protein level upon lncRAP2/*Igf2bp2* depletion in mature white adipocytes.

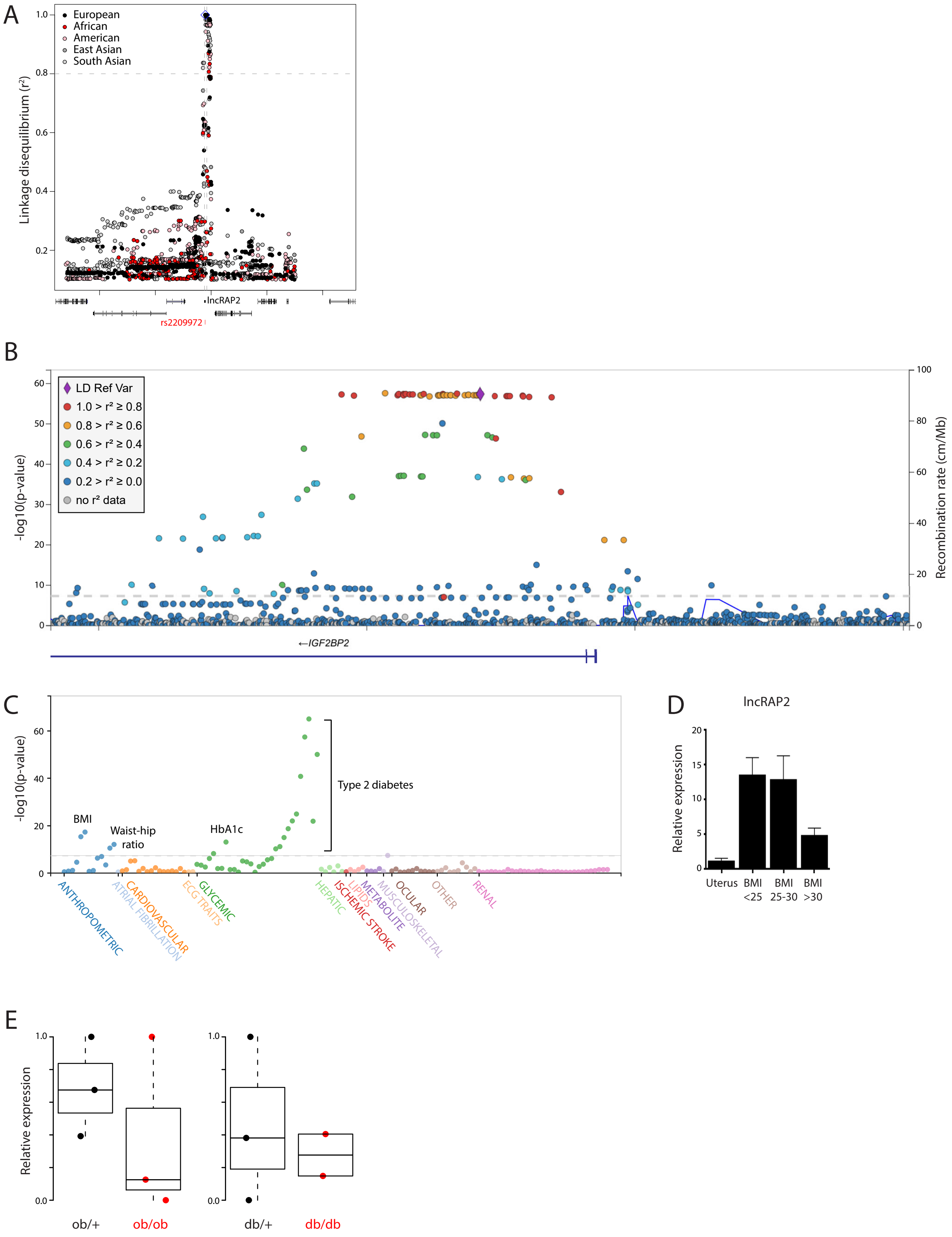
A
European
African
American
East Asian
South Asian
Linkage disequilibrium (r²)
1.0
0.8
0.6
0.4
0.2
lncRAP2
rs2209972
B
LD Ref Var
1.0 > r² ≥ 0.8
0.8 > r² ≥ 0.6
0.6 > r² ≥ 0.4
0.4 > r² ≥ 0.2
0.2 > r² ≥ 0.0
no r² data
-log10(p-value)
Recombination rate (cm/Mb)
60
50
40
30
20
10
0
100
80
60
40
20
0
←IGF2BP2
C
-log10(p-value)
60
40
20
0
BMI
Waist-hip ratio
HbA1c
Type 2 diabetes
ANTHROPOMETRIC
ATRIAL FIBRILLATION
CARDIOVASCULAR
ECG TRAITS
GLYCEMIC
HEPATIC
ISCHEMIC STROKE
LIPIDS
METABOLITE
MUSCULOSKELETAL
OCULAR
OTHER
RENAL
D
lncRAP2
Relative expression
20
15
10
5
0
Uterus
BMI <25
BMI 25-30
BMI >30
E
Relative expression
1.0
0.0
ob/+
ob/ob
1.0
0.0
db/+
db/db

**Supplementary Fig. 5 lncRAP2-Igf2bp2 genetic and expression variability are associated with obesity-linked diabetes risk.**

(**a**) Human rs2209972 is in strong linkage disequilibrium with genetic variants within lncRAP2 but no other loci within 500kb. Linkage disequilibrium with rs2209972 (diamond) based on 1000 Genomes[71] data for the indicated populations.

(**b**) Regional association with type 2 diabetes at the IGF2BP2 locus. Significance of association between genetic variants surrounding the IGF2BP2 transcription start site and type 2 diabetes in 898,130 European-descent individuals[70]. Variants are colored based on 1000 Genomes All samples[71] linkage disequilibrium with rs7615045 (diamond).

(**c**) IGF2BP2 rs7615045 is specifically associated with body mass, glycated hemoglobin (HbA1c), and type 2 diabetes. Phenome-wide association results between rs7615045 and 317 phenotypes across 16,278,030 individuals of various ancestries from the Type 2 Diabetes Knowledge Portal (http://www.type2diabetesgenetics.org/variantInfo/variantInfo/rs7615045). Only significant ($p<0.05$) associations are shown.

(**d**) Adipose lncRAP2 is progressively reduced with obesity. lncRAP2 levels in adipose tissue samples from patients with different body mass index (BMI)

(**e**) *Igf2bp2* is suppressed during diabetes onset and progression. *Igf2bp2* levels in visceral adipose tissue from leptin-deficient (ob/ob) and leptin receptor-deficient (db/db) 9-10 week-old mice[81].
